# Interhemispheric inhibition between dorsal premotor and primary motor cortices is released during preparation of unimanual but not bimanual movements

**DOI:** 10.1101/2023.07.20.549948

**Authors:** Ronan Denyer, Brian Greeley, Ian Greenhouse, Lara A. Boyd

## Abstract

Previous research applying transcranial magnetic stimulation during unimanual reaction time tasks indicates a transient change in the inhibitory influence of dorsal premotor cortex over contralateral primary motor cortex shortly after the presentation of an imperative stimulus. Interhemispheric inhibition from the dorsal premotor cortex to the contralateral primary motor cortex shifts depending on whether the targeted effector representation in the primary motor cortex is selected for movement. Further, the timing of changes in inhibition covaries with the selection demands of the reaction time task. Less is known about modulation of dorsal premotor to primary motor cortex interhemispheric inhibition during the preparation of bimanual movements. In this study, we used a dual coil transcranial magnetic stimulation to measure dorsal premotor to primary motor cortex interhemispheric inhibition between both hemispheres during unimanual and bimanual simple reaction time trials. Interhemispheric inhibition was measured early and late in the “pre-movement period” (defined as the period immediately after the onset of the imperative stimulus and before the beginning of voluntary muscle activity). We discovered that interhemispheric inhibition was more facilitatory early in the pre-movement period compared to late in the pre-movement period during unimanual reaction time trials. In contrast, interhemispheric inhibition was unchanged throughout the pre-movement period during symmetrical bimanual reaction time trials. These results suggest that there is greater interaction between the dorsal premotor cortex and contralateral primary motor cortex during preparation of unimanual actions compared to bimanual actions.

## INTRODUCTION

The dorsal premotor cortex (PMd) is a brain region critical for the translation of external information into voluntary action (Denyer et al., 2023; Picard & Strick, 2001). Evidence from non-human primate studies shows that PMd single-neuron firing rates during reach preparation correlate to kinematic features of the upcoming movement, including direction (Cisek & Kalaska, 2005; Godschalk et al., 1985; Messier & Kalaska, 2000; Riehle & Requin, 1989), trajectory (Hocherman & Wise, 1991), amplitude (Messier & Kalaska, 2000) and speed (Churchland et al., 2006). Furthermore, the application of dimensionality reduction methods to concurrent recordings of hundreds of PMd neurons has revealed distinct latent neural dynamics during reach preparation and execution, with population firing rates that are best modelled with point attractor dynamics during preparation and with rotational dynamics during execution (Churchland et al., 2010, 2012; Kaufman et al., 2014). Notably, transitions between these preparation-related and execution-related “neural states” occurs in a temporally predictable manner that correlates highly with the onset of movement (Kaufman et al., 2016; Thura et al., 2022). The finding that the activity of PMd neurons reflects kinematic features of movement at both the single-cell and population level suggests that this brain region is critical for selecting, planning, and initiating movements (Denyer et al., 2023; Duque et al., 2012).

PMd presumably influences descending motor activity through a rich network of inhibitory and facilitatory intercortical connections with the primary motor cortex (M1) (Tokuno & Nambu, 2000). Dual coil TMS can be used to non-invasively study the excitatory and inhibitory influence of PMd on M1 during motor preparation in humans at short time intervals (6-10ms) by comparing the size of motor evoked potentials (MEP) in targeted effectors in response to TMS over M1 alone (i.e., unconditioned MEP) to TMS over M1 preceded by TMS over PMd (i.e., conditioned MEP) (Mochizuki et al., 2004; Ni et al., 2009). During reaction time (RT) tasks, TMS pulses can be applied at specific timepoints before movement onset to track whether PMd increases or decreases excitability in contralateral M1 across motor preparation. During unimanual movement preparation, PMd transiently alters inhibition on cortical effector representations in contralateral M1 40-150 ms after the presentation of an imperative stimulus (i.e., the stimulus which communicates that the participant should move) (Koch et al., 2006; Kroeger et al., 2010; Liuzzi et al., 2010, 2011; O’Shea et al., 2007). Importantly, PMd-MI interhemispheric inhibition (IHI) is transiently *released* if the targeted effector is selected for movement; the net effect is to decrease inhibition (Kroeger et al., 2010; Liuzzi et al., 2010, 2011; O’Shea et al., 2007). In contrast, when the target effector is not selected for movement PMd-M1 IHI is transiently *increased* (Koch et al., 2006). Furthermore, the timing of transient changes in PMd-M1 IHI depends on the selection demands of the task. In simple RT tasks transient changes in PMd-M1 IHI occur ∼40-50 ms after the imperative cue (Liuzzi et al., 2010, 2011; O’Shea et al., 2007). During choice RT and Go/NoGo RT tasks changes in PMd-M1 IHI occur ∼75-150 ms after the imperative cue (Koch et al., 2006; Kroeger et al., 2010; O’Shea et al., 2007). These findings have led to speculation that PMd contralateral to active M1 is actively involved in shaping corticospinal output to control the selection and timing of movements during unimanual RT tasks (Denyer et al., 2023; Duque et al., 2012).

While the modulation of PMd-M1 interhemispheric circuits prior to movement onset has been well defined for unimanual movements, bimanual movements have received less attention. In one study, right-handed participants rotated circular manipulanda using both of their index fingers to move a dot along a path on a computer monitor. Modulation of left PMd-right M1 IHI depended on bimanual movement symmetry (Fujiyama et al., 2016). Specifically, when the left index finger was required to rotate 3 times faster than the right index finger, left PMd-right M1 interactions became more facilitatory 50 ms prior to the imperative stimulus. Conversely, when the left hand was required to rotate 3 times slower than the right hand, left PMd–right M1 interactions became more inhibitory. Furthermore, the degree of modulation of left PMd–right M1 IHI compared to rest correlated with lower target tracking errors in the behavioural task when asymmetric hand rotations were required. In contrast, no change in the influence of PMd on M1 was observed when both index fingers were required to move symmetrically at the same angular velocity regardless of hemisphere. This suggests that PMd-M1 interhemispheric circuits (in particular left PMd-right M1 circuits) may be important for asymmetric but not symmetric bimanual control.

How PMd-M1 interhemispheric inhibitory circuits are modulated during the pre-movement period (i.e., after the imperative stimulus but before movement onset) of bimanual RT tasks is not well understood. One study measured PMd-M1 IHI in right-handed individuals in both hemispheres during the pre-movement period and found that inhibition ratios were similar across unimanual and symmetric bimanual simple RT trials (Verstraelen et al., 2021). However, PMd-M1 IHI was assessed only at 75% of mean RT (∼110-180 ms), much later than the range in which transient switches in inhibition are observed for unimanual simple RT tasks (20% of mean RT; ∼30-40ms) (Liuzzi et al., 2010, 2011; O’Shea et al., 2007).

The current study aimed to assess how PMd-M1 IHI is modulated across early and late stages of the pre-movement period of a simple RT task, and if patterns of PMd-M1 circuit modulation depend on whether the planned response is unimanual or symmetrically bimanual. In line with previous research, we predicted that PMd-M1 IHI would be more facilitatory at 50 ms (early) compared to 100 ms (late) post imperative signal when unimanual responses are performed (**Hypothesis 1A**) (Liuzzi et al., 2010, 2011; O’Shea et al., 2007). Since evidence suggests that the preparation of symmetrical bimanual movements is not associated with significant modulation in PMd-M1 IHI 50 ms prior to the onset of the imperative cue (Fujiyama et al., 2016) or ∼40 ms prior to the onset of movement during simple RT trials (Verstraelen et al., 2021), we predicted that PMd-M1 IHI would remain unchanged across the pre-movement period when symmetrical bimanual responses are performed (**Hypothesis 1B**). PMd-M1 IHI was assessed in both hemispheres to determine whether preparation related changes in PMd-M1 IHI display hemispheric asymmetry. Since evidence suggests inhibitory output of left PMd and right PMd on contralateral M1 is similar during preparation of unimanual (Koch et al., 2006; O’Shea et al., 2007) and symmetric bimanual movements (Fujiyama et al., 2016), we further predicted that patterns of PMd-M1 IHI modulation would be similar across hemispheres (**Hypothesis 2**).

## MATERIALS AND METHODS

### Participants

Eighteen healthy individuals recruited from British Columbia completed the study. One participant was removed due to technical issues during TMS data collection. Therefore, 17 individuals (5 male, 12 female, 36.9 ± 14.5 years old) were included in the final analyses. All participants were right-handed according to the Edinburgh Handedness Inventory (Oldfield, 1971). Participant characteristics are presented in **Table 1**. All protocols were approved by Clinical Research Ethics Board at the University of British Columbia. Prior to participating, individuals were screened for contraindications to TMS and provided written informed consent in accordance with the Declaration of Helsinki.

**Table 1.** Participant characteristics. M1 – primary motor cortex; PMd – dorsal premotor cortex; RMT – resting motor threshold (percent maximum stimulator output); TPT – test pulse threshold (percent maximum stimulator output); PMd Coil refers to the 5-cm diameter figure-of-8 coil used to stimulate PMd, M1 Coil refers to the 7-cm diameter figure-of-8 coil used to stimulate M1.

| Participant | Sex | Age | Right<br>M1<sub>PMd</sub> Coil<br>RMT | Right<br>M1<sub>M1</sub> Coil<br>RMT | Right<br>M1<sub>M1</sub> Coil<br>TPT | Left<br>M1<sub>PMd</sub><br>Coil RMT | Left<br>M1<sub>M1</sub> Coil<br>RMT | Left<br>M1<sub>M1</sub> Coil<br>TPT |
| --- | --- | --- | --- | --- | --- | --- | --- | --- |
| 1 | F | 24 | 61 | 55 | 85 | 68 | 51 | 75 |
| 2 | F | 32 | 52 | 49 | 56 | 54 | 56 | 68 |
| 3 | F | 54 | 54 | 52 | 65 | 53 | 48 | 64 |
| 4 | F | 30 | 44 | 41 | 47 | 41 | 40 | 41 |
| 5 | F | 21 | 69 | 60 | 85 | 72 | 58 | 85 |
| 6 | M | 37 | 53 | 48 | 55 | 51 | 46 | 53 |
| 7 | F | 25 | 33 | 31 | 36 | 39 | 34 | 38 |
| 8 | F | 31 | 54 | 52 | 64 | 60 | 49 | 65 |
| 9 | F | 34 | 59 | 46 | 71 | 55 | 49 | 85 |
| 10 | F | 40 | 59 | 59 | 80 | 70 | 60 | 77 |
| 11 | F | 59 | 53 | 53 | 75 | 53 | 45 | 66 |
| 12 | M | 66 | 34 | 40 | 40 | 38 | 35 | 43 |
| 13 | F | 62 | 44 | 44 | 49 | 42 | 38 | 48 |
| 14 | M | 30 | 48 | 56 | 56 | 59 | 52 | 60 |
| 15 | F | 27 | 48 | 46 | 45 | 44 | 40 | 47 |
| 16 | M | 22 | 60 | 60 | 75 | 64 | 58 | 73 |
| 17 | M | 29 | 57 | 64 | 85 | 59 | 60 | 70 |

### Experimental Design

Using a within-subjects design, each participant completed two experimental sessions separated by at least 24 hours to determine whether PMd-M1 IHI is modulated across the pre-movement period of a simple RT task. Sessions differed only in terms of which hemispheres were assessed (right PMd-left M1/left PMd-right M1). The order in which hemispheres were assessed was randomized and counter-balanced between participants. During each session, PMd-M1 IHI were assessed at 50 ms and at 100 ms post an imperative visual stimulus, and also while the participant was at rest and not performing a task. During each session, participants performed 2 versions of the simple RT task: one which required unimanual responses and one which required bimanual responses (**Figure 1A**). Both versions of the task were performed in discrete blocks of 80 trials (160 total), with the order of task version counterbalanced between participants but held constant between experimental sessions. PMd- M1 IHI was measured using established dual coil TMS methods (Mochizuki et al., 2004; Ni et al., 2009).

**Figure 1.**
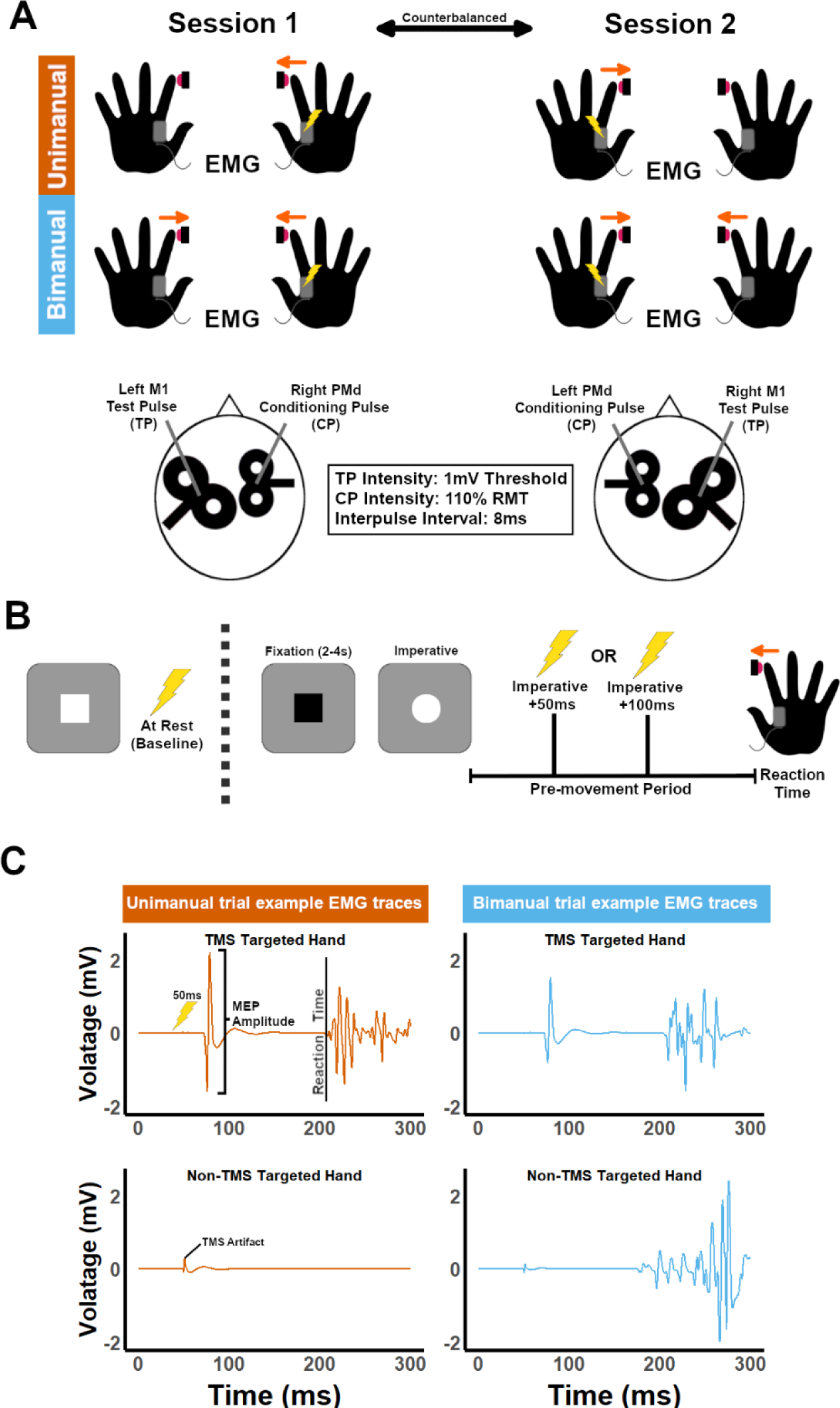
**(A)** Participants completed 2 experimental sessions separated by at least 24 hours. Sessions differed only in the hemispheric direction of PMd-M1 IHI assessed. Participants completed 80 unimanual simple reaction time (RT) trials and 80 bimanual simple RT trials during each session. Electromyography (EMG) was recorded from bilateral flexor dorsal interossei to acquire motor evoked potentials (MEPs) and infer reaction time (RT) based on EMG “bursts” on each trial. **(B)** Transcranial magnetic stimulation (TMS)/EMG measures were acquired at rest and during the pre-movement period of an SRT paradigm (lightning bolts). MEPs were elicited at either 50ms or 100ms into the pre-movement period on discrete trials. **(C)** Example EMG traces from unimanual and bimanual trials for both TMS targeted and non-TMS targeted hands for the 50ms condition. MEP amplitude and EMG RT burst onset were the primary EMG events of interest.

### Experimental Setup

Participants were seated in front of a computer monitor (BenQ, CA, USA; 144 Hz refresh rate) with both hands placed palm-down on the surface of a table (**Figure 1A**). USB-interfaced response buttons were fixed to two sides of a wooden box such that button presses could be executed starting from a resting hand position. Visual stimulus presentation was controlled by Psychtoolbox 3.0 (Kleiner et al., 2007); EMG recording and the timing of TMS stimuli was controlled using the VETA MATLAB toolbox (Jackson & Greenhouse, 2019).

### Magnetic Resonance Imaging

A 3D T1-weighted turbo field echo MRI image was used in conjunction with neuronavigation software to ensure consistent targeting of PMd and M1 by TMS across participants (TR = 7.7 milliseconds, TE = 3.5 milliseconds, flip angle θ = 8o, field of view = 256 × 256 mm, 170 slices, 1 mm slice thickness, scan time = 6.6 minutes). A Montreal Neurosciences Institute (MNI) space region of interest (ROI) mask for left and right PMd was created based on established MNI coordinates for PMd (right PMd: X=25, Y=3, Z=71; left PMd: X=-25, Y=3, Z=71) (Duque et al., 2012). T1-weighted images were non-linearly transformed into MNI space using FSL FNIRT (Jenkinson et al., 2012). The PMd ROI mask was then transformed from MNI space to native space using the inverse of the non-linear transformation warp created by FNIRT. Each individualized ROI mask was overlayed on each T1-weighted image in FSLview, and the native space coordinates for left and right PMd were visually inspected by an experimenter. Native space coordinates were later used to target PMd with TMS using Brainsight neuronavigation software (Rogue Research Inc, Montreal, QC, Canada; see **Transcranial Magnetic Stimulation**).

### Electromyography

Surface EMG was recorded using bipolar electrodes placed over both first dorsal interosseous (FDI) muscles. A ground electrode was placed over the ulnar styloid process of the right arm. EMG was sampled at 5,000 Hz, amplified x1,000, and bandpass filtered (50–450 Hz; Delsys).

### Simple Reaction Time Task

Participants completed two versions of a simple RT task; one in which unimanual responses were required and one where bimanual responses were required (**Figure 1A**). On each trial, participants were shown a fixation cue (black square) followed by an imperative cue (white circle) centrally presented on a monitor. The fixation cue was presented with a 2000-4000 ms jitter on each trial to prevent anticipatory responses (**Figure 1B**). The imperative cue disappeared after a button press or after 2000 ms if no response was registered. Participants were instructed to press the response button(s) as fast as possible following the presentation of the imperative cue and to keep their hands relaxed between button presses. EMG from left and right FDI was visually checked by the experimenters throughout each session on a second monitor situated outside of the participants’ field of vision. If resting background EMG activity was observed outside of button presses, participants were instructed to relax their hands.

During each session, participants performed a block of 80 unimanual trials and a block of 80 bimanual trials. The order of blocks was counterbalanced between participants. An opportunity for a break was provided after every 20 trials.

### Transcranial Magnetic Stimulation

#### Setup and thresholding

TMS was administered using a Magstim BiStim-200^2^ stimulator. The TMS coil used to stimulate M1 was a 7-cm diameter figure-of-8 coil, while the TMS coil used to stimulate PMd was a 5-cm diameter figure-of-8 “branding iron” coil. Brainsight neuronavigation (Rogue Research Inc, Montreal, QC, Canada) was used to ensure coil positioning was accurate and consistent across sessions. Two experimenters stood adjacent to the participant and each held one TMS coil over each region of interest. Each coil had an infrared tracker attached which allowed experimenters to track the position of the coil relative the head in real time in Brainsight throughout each session and outside of the participant’s field of vision.

For TMS over M1, the center of the coil was positioned in a posterior-to-anterior orientation 45- degrees to the mid-sagittal plane over the left/right M1 (**Figure 1A**). The “hotspot” for the FDI representation was found by probing different locations in the M1 “hand knob” area. The location at which TMS most consistently resulted in MEPs was defined as the “hotspot”. Once the M1 “hotspot” was determined, the PMd coil was placed in position over contralateral PMd, to ensure that both coils would be able to fit and sit flush on the participants head during dual-coil assessments. Small adjustments were made to the position of the M1 coil if necessary. At the “hotspot”, the resting motor threshold, defined as the lowest TMS pulse intensity that elicited five out of ten MEPs greater than or equal to a peak-to-peak amplitude of 0.05 mV, was determined. After establishing resting motor threshold, a test pulse threshold, defined as the lowest TMS pulse intensity that elicited five out of 10 MEPs greater than or equal to a peak-to-peak amplitude of 1 mV, was determined.

For TMS over PMd, the center of the coil was positioned in a lateral-to-medial orientation 90- degrees to the mid-sagittal plane (**Figure 1B**). The coordinates for PMd stimulation were defined by a standardised FSL analysis procedure applied to each participant’s T1-weighted MRI image (see **Magnetic Resonance Imaging**). Following standard procedure, PMd stimulation intensity was based off the resting motor threshold of M1 ipsilateral to PMd (Mochizuki et al., 2004; Ni et al., 2009). Therefore, the hotspot and resting motor threshold of M1 ipsilateral to the site of PMd stimulation were determined with the 5-cm PMd coil, with the coil held in posterior-to-anterior orientation 45-degrees to the mid-sagittal plane. During dual coil TMS assessments, PMd TMS intensity was set to 110% of ipsilateral M1 resting motor threshold.

To ensure M1 was not stimulated by the PMd coil during PMd-M1 IHI dual coil TMS assessments, 5 single pulses of TMS were applied over the PMd target at 110% of resting motor threshold to test whether any MEPs were elicited. If an MEP was observed, the PMd target was moved anterior by 5mm. If MEPs were still observed in 5 further trials of TMS over this adjusted target, intensity was lowered by 1% maximum stimulator output until 5 consecutive trials without MEPs were observed. PMd target/intensity adjustments were made in 4 of 17 participants, and all intensities remained suprathreshold. M1 TMS pulse intensity was set to the test pulse threshold (see **Table 1** for individual resting motor threshold and test pulse threshold values).

#### Dual coil transcranial magnetic stimulation

Dual coil TMS was used to non-invasively study the excitatory and inhibitory influence of PMd on M1. The amplitude of MEP evoked by a single pulse of TMS over M1 (i.e., unconditioned pulse) were compared to the amplitude of MEPs evoked by a pulse of TMS over M1 preceded shortly by a conditioning pulse over PMd (i.e., conditioned pulse) (Civardi et al., 2001; Mochizuki et al., 2004; Ni et al., 2009). If conditioned TMS pulses resulted in MEPs that had larger average peak-to-peak amplitude compared to MEPs elicited by unconditioned TMS pulses, then PMd had an inhibitory influence over M1. Alternatively, if conditioned TMS pulses resulted in MEPs that had larger average peak-to-peak amplitude than MEPs elicited by unconditioned pulses, then PMd had a facilitatory influence over M1.

In the current study, unconditioned MEPs were collected by delivering a test pulse (TP) over M1 alone, with stimulator intensity set to the test pulse threshold. Conditioned MEPs were collected by delivering a conditioning pulse (CP) over PMd with stimulator intensity set 110% of resting motor threshold, followed by a TP over M1, with stimulator intensity set to the test pulse threshold. The interpulse interval was set to 8 ms. These stimulation parameters have been shown to consistently elicit PMd-M1 IHI when stimulation is delivered at rest, i.e., conditioned pulses elicit MEPs of smaller amplitude than unconditioned pulses (Koch et al., 2006; Mochizuki et al., 2004; Ni et al., 2009).

Before performing the simple RT task, 20 unconditioned MEPs and 20 conditioned MEPs were collected in pseudorandomised order while participants were at rest. During these TMS trials, the participant was instructed to stare at a white fixation square and to keep their hands relaxed. These MEPs were collected to assess how PMd-M1 IHI evolved during motor preparation compared to a resting baseline.

TMS was delivered on every trial of the simple RT task, either at 50 ms (50% of trials) or 100 ms after the imperative cue (50% of trials). Equal numbers of unconditioned and conditioned MEPs were elicited at each timepoint. The timing of TMS pulses on trials was pseudorandomized so that participants could not predict their timing.

### Data Processing

Offline analysis of EMG data was performed using the VETA toolbox (Jackson & Greenhouse, 2019) and custom-automated procedures within MATLAB. EMG variables of interest included MEP peak-to peak amplitude and EMG RT burst onset (**Figure 1C**). The ‘findEMG.m’ function was used to automatically identify EMG events. All automatically identified EMG events were visually inspected by a human rater using the ‘visualizeEMG.m’ function. Adjustments were made to MEP and EMG RT burst onset and offset when necessary, using the ‘visualizeEMG.m’ function in the VETA toolbox. Any trials that contained >0.05 mV of background muscle activity in either hand 50 ms prior to the TMS artifact were removed from further analysis. Any trial in which EMG RT bursts overlapped with MEPs or vice versa were excluded from further analysis. On average, 10.4 ± 1.2% of trials were excluded per participant.

### Dependent variables

#### PMd-M1 Interhemispheric Inhibition

To measure PMd-M1 IHI, the mean peak-to-peak amplitude of 20 unconditioned MEPs (TP alone) and the mean peak-to-peak amplitude of 20 conditioned MEPs (CP + TP) were calculated for each experimental condition. The mean peak-to-peak amplitude of conditioned MEPs was then divided by the mean peak-to-peak amplitude of unconditioned MEPs to generate an PMd-M1 inhibition ratio value for each participant within each experimental condition.

#### Corticospinal Excitability

To measure corticospinal excitability, the mean peak-to-peak amplitude of 20 unconditioned MEPs (TP alone) were calculated for each participant within each experimental condition.

*Reaction Time.* To measure RT, the time difference between imperative cue onset and the onset of EMG bursts was calculated for each RT trial. The mean RT of 20 trials was then calculated for each participant within each experimental condition. For the bimanual version of the simple RT task, separate averages were calculated for each hand (the hand contralateral to M1 stimulation (i.e., TMS targeted), and the hand ipsilateral to M1 stimulation (i.e., non-TMS targeted). Separate values were calculated because the onset of an MEP in the TMS targeted hand during the pre-movement RT period typically results in a delayed RT EMG burst compared to non-TMS targeted effectors, presumably due to the onset of involuntary muscle activity and/or mechanical perturbation of the TMS targeted effector (Day et al., 1989; Hannah et al., 2018). This makes it difficult to directly compare RT values across TMS targeted and non-TMS targeted hands.

### Statistical Analysis

All statistical analyses were carried out in RStudio. *Post hoc* analyses were performed using Bonferroni’s correction for multiple comparisons where appropriate.

#### Primary analyses

*PMd-M1 Interhemispheric Inhibition.* To test if PMd-M1 IHI changes differently across time during preparation of unimanual and bimanual movements (**Hypothesis 1**), and whether changes in PMd-M1 IHI depend on the hemispheres assessed (**Hypothesis 2**), a three-way repeated measures analysis of variance (RM-ANOVA) was performed using within-subjects factors STIMULUS ONSET ASYNCHRONY (50 ms, 100 ms), TASK VERSION (UNIMANUAL, BIMANUAL), and HEMISPHERE (RIGHT PMd/LEFT M1, LEFT PMd/RIGHT M1). Measures of PMd-M1 IHI recorded at REST were included in post hoc analyses to assess how PMd-M1 IHI values changed during motor preparation compared to a resting baseline.

#### Supplementary analyses

##### Corticospinal Excitability

To assess whether hypothesized changes in PMd-M1 IHI described by Hypotheses 1 and 2 were driven by changes in corticospinal excitability, a three-way RM-ANOVA was performed using within-subjects factors STIMULUS ONSET ASYNCHRONY (50 ms, 100 ms), TASK VERSION (UNIMANUAL, BIMANUAL), and HEMISPHERE (LEFT M1, RIGHT M1). Measures of corticospinal excitability recorded at rest were included in *post hoc* analyses to assess how corticospinal excitability values changed during motor preparation compared to a resting baseline.

##### Reaction time

To assess whether hypothesized changes in PMd-M1 IHI described by Hypotheses 1 and 2 were driven by differences in RT across conditions of interest, two RM- ANOVAs were performed for TMS targeted and non-TMS targeted hands. For the TMS targeted hand, a four-way RM-ANOVA was performed using within-subjects factors STIMULUS ONSET ASYNCHRONY (50 ms, 100 ms), TASK VERSION (UNIMANUAL, BIMANUAL), HEMISPHERE (LEFT M1, RIGHT M1), and TMS TYPE (UNCONDITIONED, CONDITIONED). For the non-

TMS targeted hand, a three-way RM-ANOVA was performed using within-subjects factors (50 ms, 100 ms), HEMISPHERE (LEFT M1, RIGHT M1), and TMS TYPE (UNCONDITIONED, CONDITIONED).

#### Exploratory analyses

*Associations between PMd-M1 Interhemispheric Inhibition and Reaction Time.* Based on results from *a priori* planned RM-ANOVAs to test Hypothesis 1 and 2, an exploratory *a posteriori* analysis was performed to examine the effect of individual differences in RT on PMd-M1 IHI across stimulus onset asynchronies, task version, and hemisphere conditions. To test for the possible effect of mean RT on PMd-M1 IHI across factors of interest, a three-way repeated measures analysis of covariance (RM-ANCOVA) was performed using within-subjects factors STIMULUS ONSET ASYNCHRONY (50 ms, 100 ms), TASK VERSION (UNIMANUAL, BIMANUAL), and HEMISPHERE (LEFT M1, RIGHT M1), and covariate MEAN RT. Since no true “non-stimulation” RT trials were collected in this experiment, the mean of all non-stimulated hand RT values recorded on bimanual trials was used as a proxy for non-stimulation RT (Day et al., 1989; Hannah et al., 2018). Exploratory *post hoc* bivariate Pearson correlations were performed to assess the direction and strength of the relationship between PMd-M1-IHI and non-stimulated mean RT across significant within-subject factors.

## RESULTS

### PMd-M1 Interhemispheric Inhibition

For PMd-M1 IHI, a significant STIMULUS ONSET AYNCHRONY x TASK VERSION interaction effect was found (*F*1, 16 = 5.21, *p* < 0.05, η_p_ ^2^ = 0.25). To interrogate the interaction effect, the data was split by TASK VERSION and 2 follow-up two-way ANOVAs with within-subject factors STIMULUS ONSET AYNCHRONY and HEMISPHERE were run. PMd-M1 IHI data collected for each HEMISPERE at rest was added in as an extra level of STIMULUS ONSET ASYNCHRONY in both follow up ANOVAs. No significant effects were found for the bimanual task condition. For the unimanual task condition, a significant main effect of STIMULUS ONSET ASYNCHRONY was found (*F*2, 32 = 4.51, *p* < 0.05, η_p_ ^2^ = 0.22). To understand the main effect of STIMULUS ONSET ASYNCHRONY found for the unimanual task condition, pairwise t-tests were run with data pooled across HEMISPHERE. PMd-M1 IHI was significantly greater at 50 ms compared to rest (*t*(16) = 3.08, *p* < 0.05; 50 ms: 1.03 ± 0.15, Rest: 0.91 ± 0.14), and PMd-M1 IHI was significantly lower at 100 ms compared to 50 ms, however this effect did not survive corrections for multiple comparisons rest (*t*(16) = 2.42, *p* = 0.08; 100ms: 0.92 ± 0.09) (**Figure 2**).

**Figure 2.**
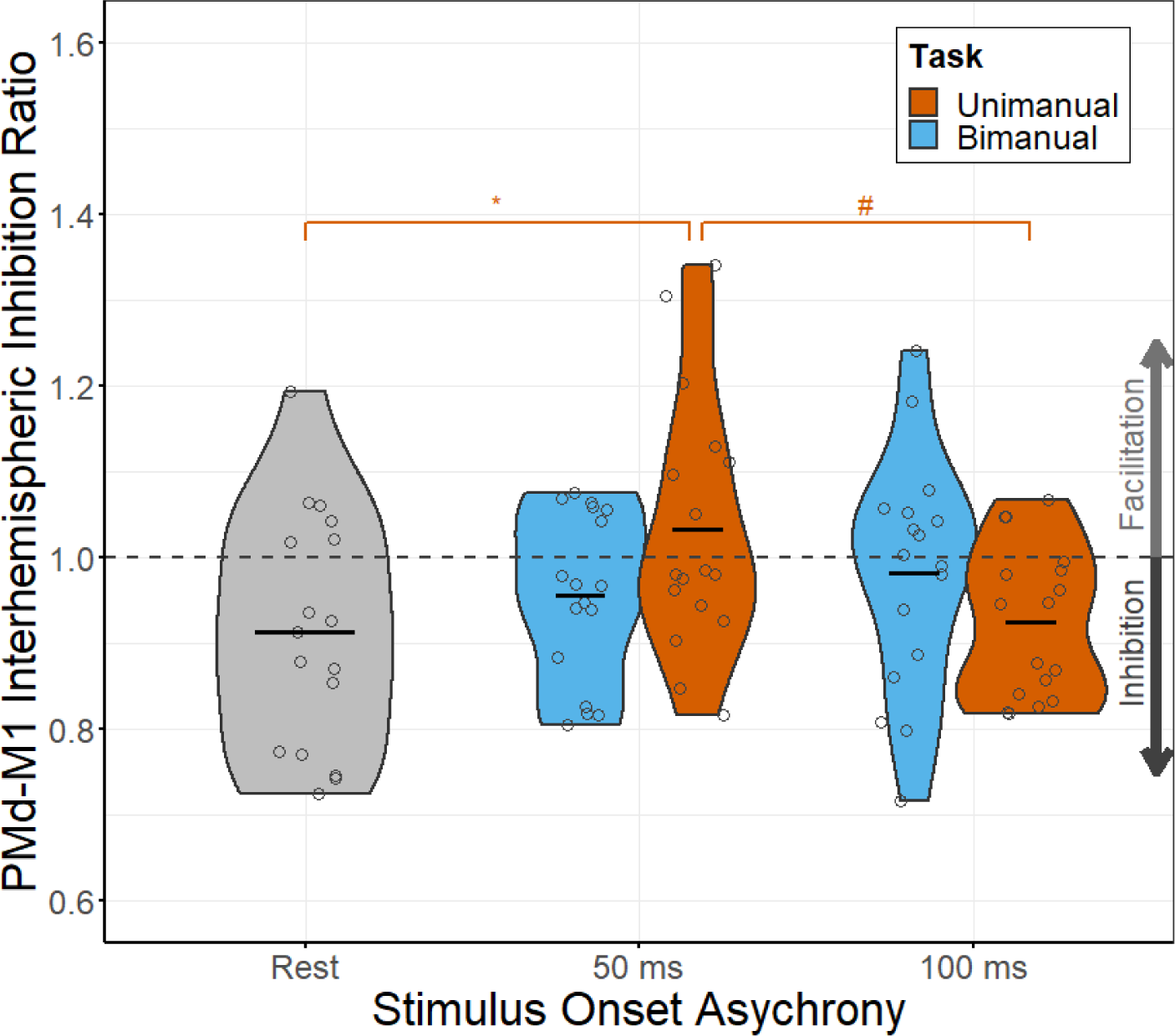
Dorsal premotor cortex (PMd) - primary motor cortex (M1) interhemispheric inhibition as assessed by dual coil transcranial magnetic stimulation at rest and across preparation of unimanual and bimanual movements. Data are pooled across hemisphere. PMd-M1 interhemispheric inhibition ratios switched from an inhibition resting state to facilitation 50ms post imperative stimulus and back to inhibition at 100ms post imperative stimulus during unimanual trials. PMd-M1 interhemispheric inhibition ratios remained inhibitory during bimanual trials. Black horizontal lines represent the mean. For unimanual task comparisons, # *p* < 0.05 before corrections for multiple comparisons, \**p* < 0.05 after corrections for multiple comparisons.

### Corticospinal Excitability

For corticospinal excitability, a main effect of STIMULUS ONSET AYNCHRONY (*F*1, 16 = 9.38, *p* < 0.01, η_p_ ^2^ = 0.37) was found. This main effect should be considered in the context of a significant STIMULUS ONSET AYNCHRONY x HEMISPHERE interaction effect (*F*1, 16 = 6.18, *p* < 0.05, η_p_ ^2^ = 0.28). To interrogate the interaction effect, the data was pooled across TASK VERSION, split by HEMISPHERE, and follow-up pairwise t-tests across the factor of STIMULUS ONSET AYNCHRONY were performed. Measures of left M1 and right M1 corticospinal excitability recorded at rest were included as a level of STIMULUS ONSET ASYNCHRONY to further assess how excitability changed at 50 ms and 100 ms compared to rest for each hemisphere. For left M1, corticospinal excitability was significantly greater at 100 ms compared to 50 ms (*t*(16) = 3.56, *p* < 0.01; 50 ms: 2.99 ± 1.37mV, 100 ms: 3.41 ± 1.52mV). Left M1 corticospinal excitability was also significantly greater at 50ms (*t*(16) = 6.22, *p* < 0.0001; Rest: 1.35 ± 0.61mV) and at 100 ms (*t*(16) = 7.2, *p* < 0.0001) compared to rest (**Figure 3A**). For right M1, there was no significant difference at 100 ms compared to 50 ms. However, right M1 corticospinal excitability was significantly greater at 50 ms (*t*(16) = 4.58, *p* < 0.001; Rest: 1.3 ± 0.49mV, 50 ms: 2.48 ± 1.11mV), and at 100ms (*t*(16) = 4.47, *p* < 0.005; 100 ms: 2.62 ± 1.28mV) compared to at rest. In summary, corticospinal excitability increased significantly during preparation of unimanual and bimanual responses compared to rest for left and right M1, and was significantly greater at 100ms compared to 50ms stimulus onset asynchrony for left M1 (**Figure 3B**).

**Figure 3.**
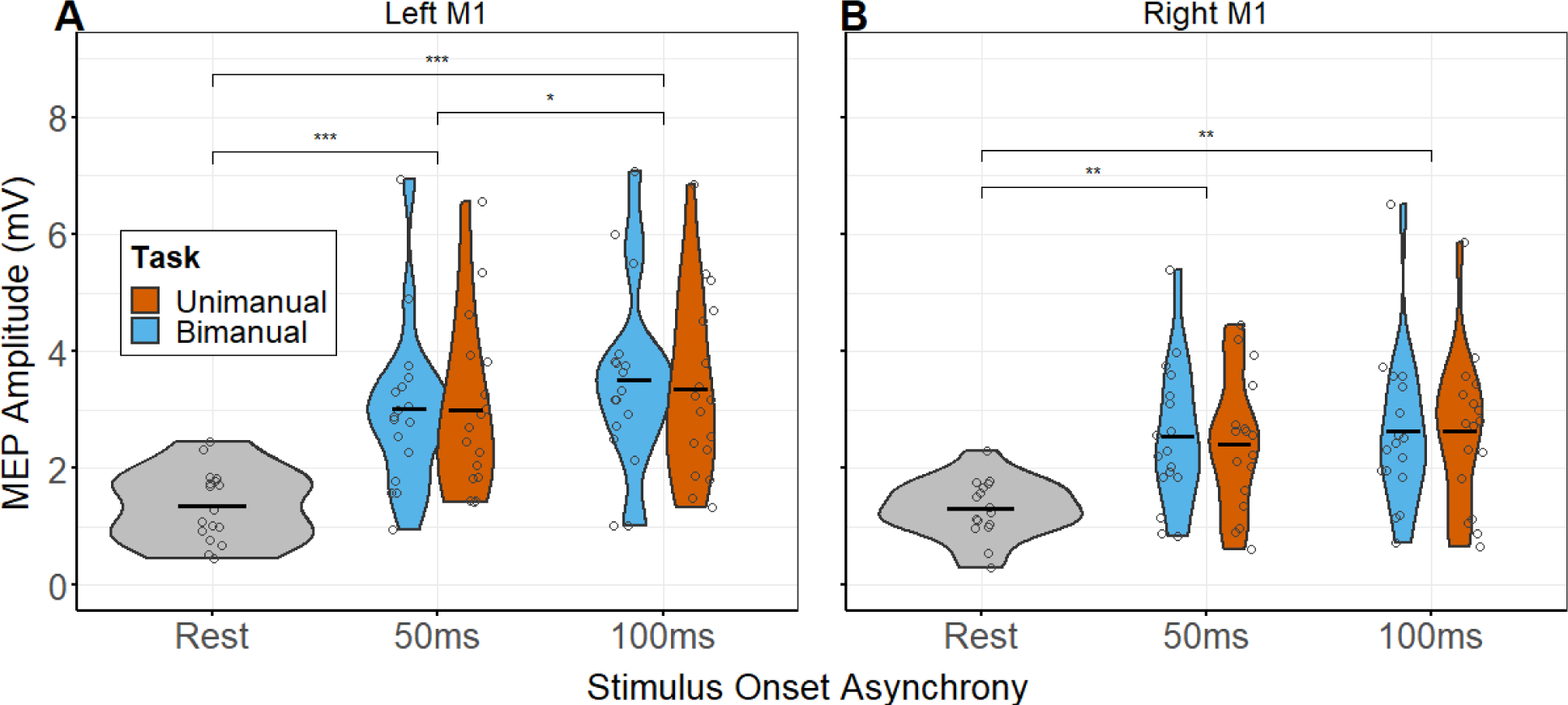
Corticospinal excitability as measured by peak-to-peak motor evoked potentials (MEPs) assessed by single pulse transcranial magnetic stimulation in left **(A)** and right **(B)** primary motor cortex (M1). Both left and right M1 showed significant increases in corticospinal excitability when recorded at 50 and 100ms post imperative stimulus compared to at rest. Only left M1 showed an increase in corticospinal excitability at 100ms compared to 50ms post imperative stimulus. Changes in corticospinal excitability were similar across unimanual and bimanual trials. Black horizontal lines represent the mean, and dots represent individual participants. \**p <* 0.01, \*\**p* < 0.005, \*\*\**p* < 0.0001.

### Reaction Time

For TMS targeted hand RT, main effects of STIMULUS ONSET AYNCHRONY (*F*1, 16 = 368.76, *p* < 0.0001, η_p_ ^2^ = 0.96; 50 ms trials: 213 ± 29 ms, 100ms trials: 249 ± 34 ms; **Figure 4A**), TMS TYPE (*F*1, 16 = 28.2, *p* < 0.0001, η_p_ ^2^ = 0.64; conditioned TMS trials: 228 ± 37 ms, unconditioned TMS trials: 235 ± 35 ms), and TASK VERSION (*F*1, 16 = 4.6, *p* < 0.05, η_p_ ^2^ = 0.22; unimanual trials: 229 ± 37 ms, bimanual trials: 234 ± 35 ms) were found. TMS TYPE and TASK VERSION main effects should be interpreted in the context of a significant HEMISPHERE x TMS TYPE x TASK VERSION interaction effect (*F*1, 16 = 11.54, *p* < 0.01, η_p_ ^2^ = 0.42). To further assess this three-way interaction, data were pooled across stimulus onset asynchronies and split by task version. Two separate two-way RM-ANOVAs were then performed using within subject-factors HEMISPHERE, and TMS TYPE. For unimanual trials, a main effect of TMS TYPE (*F*1, 16 = 17.2, *p* < 0.001, η_p_ ^2^ = 0.52; conditioned TMS trials: 225 ± 38 ms, unconditioned TMS trials: 231 ± 36 ms) was found (**Figure 4B**). For bimanual trials, a main effect of TMS TYPE (*F*1, 16 = 15.13, *p* < 0.01, η_p_ ^2^ = 0.49, conditioned TMS trials: 230 ± 29 ms, unconditioned TMS trials: 237 ± 29 ms) was also found, as well as a HEMISPHERE x TMS TYPE interaction effect (*F*1, 16 = 20.89, *p* < 0.001, η_p_ ^2^ = 0.57). To assess this interaction effect, bimanual task data were split by HEMISPHERE, and two pairwise t-tests were performed with within-subject factor TMS TYPE. For bimanual trials in which left M1 was stimulated, there was a no difference in RT across unconditioned TMS and conditioned TMS trials. For bimanual trials in which right M1 was stimulated RT was significantly lower on conditioned TMS bimanual trials compared to unconditioned TMS bimanual trials (*t*(16) = 5.35, *p* > 0.0001; conditioned TMS trials: 226 ± 25 ms, unconditioned TMS trials: 238 ± 26 ms) (**Figure 4B**).

**Figure 4.**
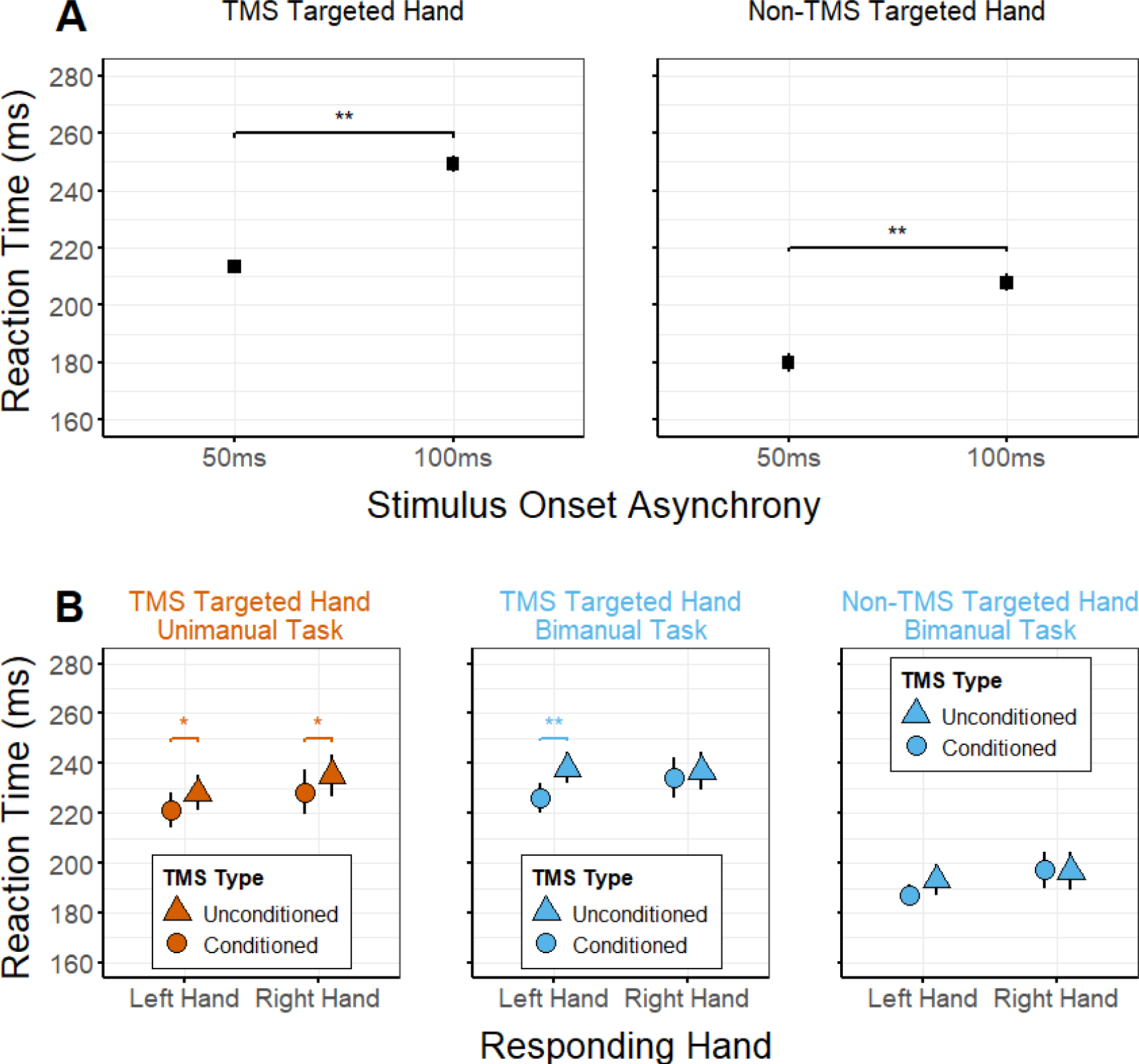
**(A)** Reaction time for each stimulus onset asynchrony for TMS targeted (left) and non-TMS targeted (right; bimanual trials only) hands. Data pooled over other within subject factors of hemisphere, TMS pulse type and task version (unimanual trials only – non-TMS targeted hand RT was not collected on unimanual trials). **(B)** Reaction time for left and right hands and single and double TMS pulse types, shown for unimanual trials (left), the TMS targeted hand on bimanual trials (middle), and the non-TMS targeted hand on bimanual trials (right). RT was defined as the time between the onset of the imperative stimulus and the onset of EMG burst activity. \**p*<0.001, \*\**p*<0.0001.

For non-TMS targeted hand RT, a main effect of STIMULUS ONSET AYNCHRONY (*F*1, 16 = 448.32, *p* < 0.0001, η_p_ ^2^= 0.97; 50ms trials: 180 ± 29 ms, 100 ms trials: 208 ± 26 ms; **Figure 4A**) and a HEMISPHERE x TMS TYPE interaction (*F*1, 16 = 8.9, *p* < 0.01, η_p_ ^2^ = 0.36) were found. To further assess the interaction effect, data were pooled across stimulus onset asynchronies and split by hemisphere. Follow-up pairwise t-tests for each hemisphere found a non-significant trend for faster RT on conditioned TMS bimanual trials compared to unconditioned TMS bimanual trials for the left PMd/right M1 configuration (t(16) = 2.03, *p* = 0.059; conditioned TMS trials: 187 ± 23 ms, unconditioned TMS trials: 193 ± 30 ms) but not for the right PMd/left M1 configuration (t(16) = 0.21, *p* = 0.835; conditioned TMS trials: 198 ± 34 ms, unconditioned TMS trials: 197 ± 33 ms; **Figure 4B**).

### Associations between PMd-M1 Interhemispheric Inhibition and Reaction Time

A main effect of STIMULUS ONSET ASYNCHRONY was found (*F*1, 16 = 5.9, *p* < 0.05, η_p_^2^ = 0.28). Furthermore, STIMULUS ONSET ASYNCHRONY x MEAN RT (*F*1, 16 = 10.46, *p* < 0.01, η_p_ ^2^ = 0.41), HEMISPHERE x MEAN RT (*F*1, 15 = 5.37, *p* < 0.05, η_p_ ^2^ = 0.26), and STIMULUS ONSET ASYNCHRONY x TASK VERSION interaction effects were also found (*F*1, 15 = 5.73, *p* < 0.05, η_p_ ^2^ = 0.28). To further unpack the STIMULUS ONSET ASYNCHRONY x TASK VERSION and STIMULUS ONSET ASYNCHRONY x MEAN RT interaction effects, the data were split by TASK VERSION and two follow up RM-ANCOVAs with within-subject factors STIMULUS ONSET AYNCHRONY and HEMISPHERE were run with MEAN RT included as a covariate. For the unimanual task version, a main effect of STIMULUS ONSET AYNCHRONY (*F*1, 15 = 8.33, *p* < 0.05, η_p_ ^2^ = 0.36) and a STIMULUS ONSET AYNCHRONY x MEAN RT (*F*1, 15 = 6.72 *p* < 0.05, η_p_^2^ = 0.31) interaction effect were found. There were no significant effects for the bimanual task version.

To further interrogate the effect of stimulus onset asynchrony on the strength and direction of the relationship between PMd-M1 IHI and mean RT in the unimanual task condition, Pearson bivariate correlations were performed between measures of PMd-M1 IHI collected at each STIMULUS ONSET ASYNCHRONY and MEAN RT. MEAN RT was expressed as a percentage of STIMULUS ONSET ASYNCHRONY to aid with interpretation. PMd-M1 IHI was shown to significantly correlate with MEAN RT when measured at 50 ms post imperative stimulus (ρ = - 0.4, *p* < 0.05; **Figure 5**) but not at 100 ms post imperative stimulus (ρ = 0.212, *p* = 0.23).

**Figure 5.**
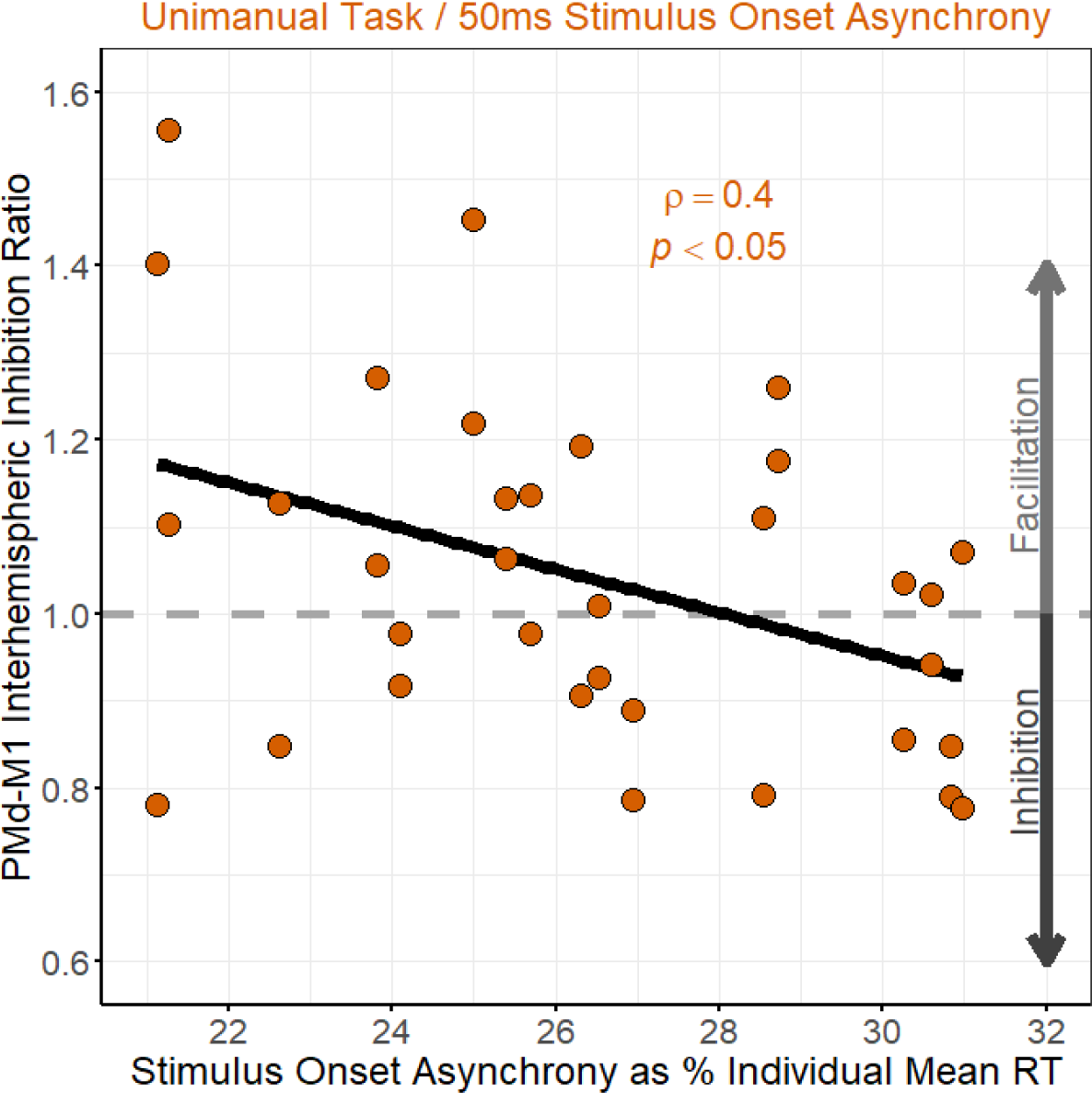
Correlation between mean RT and PMd-M1 IHI recorded during unimanual trials at a stimulus onset of asynchrony of 50ms. Facilitation tended to be observed when 50ms represented 20-28% of an individual’s mean RT, while inhibition was more likely when 50ms represented >28% of an individual’s mean RT.

## DISCUSSION

In the current study, we investigated the modulation of PMd-M1 IHI across two timepoints within the pre-movement period (i.e., the period of time between the onset of the imperative stimulus and the onset of movement) of a visually cued unimanual and symmetrically bimanual simple RT task. For unimanual movements, we found that PMd-M1 interhemispheric interactions transiently switched from inhibitory to facilitatory early (50 ms) and back to inhibitory later (100 ms) within the pre-movement period (**Figure 2**). Additionally, the strength of PMd-M1 IHI at 50 ms correlated with mean RT (**Figure 5**). In contrast, PMd-M1 interhemispheric interactions remained inhibitory throughout the pre-movement period when bimanual movements were required (**Figure 2**), and were unrelated to RT. Taken together, these results suggest that discrete unimanual and bimanual movements may be preceded by different patterns of excitability in PMd-M1 interhemispheric circuits.

### PMd-M1 Interhemispheric Interactions Differ for Unimanual and Bimanual Movements

During the pre-movement period of unimanual movements, PMd-M1 IHI shifted to interhemispheric facilitation 50 ms post imperative stimulus before returning to IHI 100 ms post imperative stimulus across both hemispheres (**Figure 2**). These findings are consistent with previous investigations that showed PMd-M1 interhemispheric circuits in both hemispheres are involved early in the pre-movement period of unimanual movements (Koch et al., 2006; Kroeger et al., 2010; Liuzzi et al., 2010, 2011; O’Shea et al., 2007). Similar effects are noted across a variety of externally cued RT tasks, including simple RT (Liuzzi et al., 2010, 2011; O’Shea et al., 2007), choice RT (Koch et al., 2006; O’Shea et al., 2007), and Go/NoGo RT (Kroeger et al., 2010) tasks. Importantly, this research has demonstrated that switches from inhibition to facilitation are not sustained throughout the pre-movement period but are transient, occurring in a time window of ∼25 ms. Furthermore, the timing of transient switches in PMd-M1 IHI within the pre-movement period depends on the selection demands of the task, with transient switches occurring ∼40-50 ms after the imperative cue in simple RT tasks with fast mean RTs (∼200 ms) (Liuzzi et al., 2010, 2011; O’Shea et al., 2007) and at ∼75-150 ms after the imperative cue in choice RT tasks with relatively slower mean RTs (∼350 ms) (Koch et al., 2006; Kroeger et al., 2010; O’Shea et al., 2007).

While there was a significant effect of stimulus onset asynchrony on PMd-M1 IHI ratios for unimanual movements, our mean IHI ratio of 1.05 at 50 ms was lower relative to other studies using a similar paradigm, where mean ratios of 1.15-1.25 were reported (Liuzzi et al., 2010, 2011; O’Shea et al., 2007). Past work individualized the timing of TMS stimuli to 20% of each participants’ mean RT rather than using a fixed stimulus onset asynchrony (as was the case in the current study) and showed a larger average PMd-M1 IHI ratio of 1.25 at this timepoint (Liuzzi et al., 2010). To assess whether individual differences in average RT were driving the evolution of PMd-M1 IHI across unimanual preparation, we performed an additional analysis with the same within-subjects factors but included mean RT as a covariate. We observed that the strength of PMd-M1 facilitation at 50 ms post imperative stimulus correlated negatively with RT when 50 ms was expressed as a percentage of individual mean RT (**Figure 5**). When the 50 ms fixed stimulus onset asynchrony equated to 20-28% of the participants mean RT, IHI ratios tended to be facilitatory; however, when 50 ms equated to >28% of mean RT, IHI ratios tended to be inhibitory. This percentage of RT may represent a rough temporal threshold that demarcates “early” vs “late” phases of the pre-movement period, after which PMd facilitatory input to the selected effector representation in contralateral M1 is diminished. Given that 50 ms represented >28% of mean RT for a subset of participants in our study (6/17), the use of fixed TMS pulse timing of 50 ms may not have occurred early enough within the pre-movement period to identify a switch from inhibition towards facilitation in these participants. This may explain the discrepancy between the mean PMd-M1 IHI ratio reported in the current study and previous investigations, and further supports the notion that switches in PMd-M1 interhemispheric circuits are transient and important for the timing of unimanual discrete movements (O’Shea et al., 2007).

PMd-M1 IHI remained unchanged throughout the pre-movement period for bimanual movements. The absence of an effect of stimulus onset asynchrony did not appear to be due to individual differences in mean RT, since mean RT was not a significant factor for bimanual trials in the exploratory analysis. This result is in line with previous work showing that PMd-M1 IHI does not differ from baseline levels during motor preparation 50 ms before the presentation of an imperative stimulus of a bimanual task where symmetrical circular finger movements are required (Fujiyama et al., 2016). It also aligns with other work showing PMd-M1 interhemispheric interactions do not change significantly from baseline values during late stages of the pre-movement period of a simple RT task (75% of mean RT) (Verstraelen et al., 2021).

The current study adds to this body of literature, demonstrating that PMd-M1 interactions remain inhibitory early (50 ms) in the pre-movement period of simple bimanual RT trials. Taken together, these findings suggests that even simple discrete symmetrical bimanual finger movements may be preceded by fundamentally different patterns of activity between PMd and M1 in the contralateral hemisphere.

### Changes in Corticospinal Excitability are Similar for Unimanual and Bimanual Movements

Unlike PMd-M1 interhemispheric interactions, modulation of corticospinal excitability was similar across preparation of bimanual and unimanual movements. For both task versions, there was a large increase in corticospinal excitability in the pre-movement period relative to rest. Change in corticospinal excitability across the pre-movement period differed between hemispheres, with significantly larger MEP amplitudes observed at 100 ms post imperative stimulus compared to 50 ms post imperative stimulus when left M1 was stimulated but not when right M1 was stimulated (**Figure 3**). While an effect of hemisphere on corticospinal excitability was not anticipated, this finding could be attributed to differences in average RT between dominant and non-dominant hands. While we did not find a significant difference in RT between right and left index fingers on unimanual RT trials, the onset of MEPs instituted a broad slowing of RTs compared to non-TMS targeted hand RTs (**Figure 4**), making RT data difficult to interpret. It is well established that RT is slower and more variable in the non-dominant hand versus the dominant hand (Serrien et al., 2012; Shen & Franz, 2005; Taniguchi et al., 2001). Previous investigations showed that corticospinal excitability tends to remain stable or become reduced through the early stages of the pre-movement period and begin to rapidly increase ∼60 ms prior to movement onset (Duque et al., 2014; Greenhouse et al., 2015; Klein et al., 2016; Leocani et al., 2000). Given that all participants were right hand dominant, the stimulus onset asynchrony of 100 ms was likely closer to the onset of voluntary muscle activity in the right index finger when left M1 was stimulated, which might explain why corticospinal excitability increased between 50 ms and 100 ms stimulus onset asynchronies in left M1.

The difference between rest and pre-movement period corticospinal excitability may have been context dependent. Resting single pulse MEPs were collected with the participant staring at a fixation square and not planning to make any movements. Previous research shows that there is a broad increase in corticospinal excitability of the selected effector during simple RT tasks when participants are certain of the identity of an upcoming movement, possibly due to modulation of attentional resources (Mars et al., 2007; Wright et al., 2018). Therefore, as participants prepared to perform RT trials in the current study, there may have been an attentional shift which increased baseline corticospinal excitability. Future studies should account for this general increase in corticospinal excitability by thresholding M1 TMS test pulse intensity during performance of RT trials and by collecting MEPs during task performance outside of the pre-movement period, such as when the preparatory signal is first displayed.

Alternatively, a delayed response task paradigm with a pre-cued preparation period could be implemented to limit attention to a particular response during inter trial rest periods. Overall, the corticospinal excitability results suggest that the task specific effects we describe for PMd-M1 IHI are constrained to differences in activity in PMd-M1 interhemispheric circuits and not driven by differences in corticospinal excitability.

### The Functional Role of PMd-M1 Interhemispheric Circuits

Although counterintuitive, electrocorticography (Bundy et al., 2018; Ganguly et al., 2009; Merrick et al., 2022) and functional MRI (Diedrichsen et al., 2013) studies provide evidence that the PMd ipsilateral to the moving hand plays a role in unimanual movements. Furthermore, patients with unilateral M1 lesions exhibit motor control deficits in the less affected hand (Noskin et al., 2008) and skilled unimanual sequence production is impaired in healthy individuals after inhibiting the ipsilateral M1 with repetitive TMS (Chen et al., 1997). Thus, it appears that the ipsilateral cortex plays a role in shaping unimanual movement.

In dual coil TMS studies, the timing of inhibitory and facilitatory switches in PMd-M1 IHI correlate with the onset of voluntary muscle activity (O’Shea et al., 2007), and the direction of inhibition (i.e., increasing or releasing) covaries with whether an effector is selected for an upcoming movement (Koch et al., 2006). Intracranial recordings suggest this is not an inhibitory gating mechanism (for review see Denyer et al., 2023). Perhaps the transient alterations in PMd-M1 IHI during preparation instead reflect the transition of neural population responses from output-null to output-potent “neural spaces” necessary for movement execution according to the dynamical systems framework for movement preparation (Kaufman et al., 2014; Sussillo et al., 2015). Previous work demonstrates that the timing of such neural space transitions correlates with movement onset on a trial-to-trial basis and occur ∼150 ms before movement onset (Kaufman et al., 2016). In the current study, transient switches in PMd-M1 IHI during unimanual movements were stronger when sampled ∼150 ms prior to movement onset (**Figure 5**), and previous studies further suggest the timing of transient switches in PMd-M1 interactions in the pre-movement period may correlate with RT (for a review see Denyer et al., 2023). Work that combines TMS with single neuron recordings may be required to clarify this question. However, the overall evidence suggests that activity in PMd-M1 IHI circuits may be important for accurate timing of unimanual movements.

If the contralateral PMd is necessary for accurate timing of unimanual movements, then how might symmetrical bimanual movements be initiated in the absence of this input? One possibility is that cortical activity is separated during symmetrical bimanual movements but not during unimanual movements. In this scenario, PMd of each hemisphere interacts with ipsilateral but not contralateral M1. Previous TMS research has shown that the balance of inhibition within PMd-M1 intrahemispheric circuits is modulated in a similar way to PMd-M1 interhemispheric circuits during a unimanual choice RT task (Groppa, Schlaak, et al., 2012; Groppa, Werner-Petroll, et al., 2012). It may be the case that PMd-M1 intrahemispheric circuits show a similar pattern of modulation during the initiation of discrete symmetrical bimanual movements, providing a mechanism for movement initiation in the absence of input from contralateral PMd. Another possibility is that other cortical regions are selectively engaged during planning and execution of bimanual movements. The supplementary motor area (SMA) has been hypothesized to play a specialized role in bimanual behaviours (Brinkman, 1984; Kazennikov et al., 1998; Wiesendanger et al., 1994), and dual-coil TMS protocols have been used to measure the inhibitory influence of SMA on M1 (Arai et al., 2012; Rurak et al., 2021a). Indeed, it has been shown that larger SMA-M1 inhibition ratios recorded at rest correlate with greater bimanual but not unimanual task performance (Rurak et al., 2021b), suggesting a potential specialized role for this circuit in bimanual control. The application of dual coil TMS methods at different stimulus onset asynchronies during performance of bimanual RT trials could help elucidate whether modulation of inhibition through SMA-M1 circuits during preparation shows a similar pattern to that observed in PMd-M1 interhemispheric circuits for unimanual trials.

## CONCLUSION

We observed that PMd-M1 IHI shows a different pattern of change during preparation for unimanual vs bimanual movements. Our data suggest that even simple unimanual and bimanual actions are preceded by distinct patterns of change in excitability within human interhemispheric circuits. Previous work hinted at hemispheric asymmetry in the modulation of PMd-M1 interhemispheric circuits during action preparation (Koch et al., 2006; O’Shea et al., 2007). However, we found no significant difference between hemispheres across unimanual or bimanual action preparation. The current study may have clinical relevance, in particular for individuals who experienced a stroke. MRI studies demonstrate that the structural integrity of interhemispheric pathways connecting premotor and prefrontal cortices correlates with motor impairment in chronic stroke (Hayward et al., 2017) and that the functional connectivity between contralesional PMd and ipsilesional M1 correlates with motor recovery after stroke (Bestmann et al., 2010). Furthermore, recent non-human primate research demonstrate that the blockage of contralesional PMd with chemogenetics impaired dexterous hand movements in the acute recovery stage after stroke (Mitsuhashi et al., 2023). Therefore, applying the TMS techniques used in the current study to investigate PMd-M1 circuits across movement preparation in stroke populations may be a fruitful avenue to further inform basic questions about the role of PMd in motor control and also improve understanding of the neural basis of recovery after stroke.

## Data Availability Statement

The data that support the findings of this study and associated analysis code are openly available at Open Science Framework: http://doi.org/10.17605/OSF.IO/TU6N2.

## Acknowledgments

Support was provided to RD by the Canadian Partnership in Stroke Recovery. Funding for this work was provided by the Natural Sciences and Engineering Research Council of Canada (RGPIN-2017-04154, Principal Investigator: LAB).

## References

Arai, N., Lu, M.-K., Ugawa, Y., & Ziemann, U. (2012). Effective connectivity between human supplementary motor area and primary motor cortex: A paired-coil TMS study. Experimental Brain Research, 220(1), 79–87. https://doi.org/10.1007/s00221-012-3117-5

Bestmann, S., Swayne, O., Blankenburg, F., Ruff, C. C., Teo, J., Weiskopf, N., Driver, J., Rothwell, J. C., & Ward, N. S. (2010). The role of contralesional dorsal premotor cortex after stroke as studied with concurrent TMS-fMRI. J Neurosci, 30(36), 11926–11937. https://doi.org/10.1523/JNEUROSCI.5642-09.2010

Brinkman, C. (1984). Supplementary motor area of the monkey’s cerebral cortex: Short- and long-term deficits after unilateral ablation and the effects of subsequent callosal section. The Journal of Neuroscience: The Official Journal of the Society for Neuroscience, 4(4), 918–929. https://doi.org/10.1523/JNEUROSCI.04-04-00918.1984

Bundy, D. T., Szrama, N., Pahwa, M., & Leuthardt, E. C. (2018). Unilateral, 3D Arm Movement Kinematics Are Encoded in Ipsilateral Human Cortex. Journal of Neuroscience, 38(47), 10042–10056. https://doi.org/10.1523/JNEUROSCI.0015-18.2018

Chen, R., Gerloff, C., Hallett, M., & Cohen, L. G. (1997). Involvement of the ipsilateral motor cortex in finger movements of different complexities. Annals of Neurology, 41(2), 247–254. https://doi.org/10.1002/ana.410410216

Churchland, M. M., Cunningham, J. P., Kaufman, M. T., Foster, J. D., Nuyujukian, P., Ryu, S. I., & Shenoy, K. V. (2012). Neural population dynamics during reaching. Nature, 487(7405), Article 7405. https://doi.org/10.1038/nature11129

Churchland, M. M., Cunningham, J. P., Kaufman, M. T., Ryu, S. I., & Shenoy, K. V. (2010). Cortical preparatory activity: Representation of movement or first cog in a dynamical machine? Neuron, 68(3), 387–400.

Churchland, M. M., Santhanam, G., & Shenoy, K. V. (2006). Preparatory activity in premotor and motor cortex reflects the speed of the upcoming reach. J Neurophysiol, 96(6), 3130– 3146. https://doi.org/10.1152/jn.00307.2006

Cisek, P., & Kalaska, J. F. (2005). Neural correlates of reaching decisions in dorsal premotor cortex: Specification of multiple direction choices and final selection of action. Neuron, 45(5), 801–814.

Day, B. L., Rothwell, J. C., Thompson, P. D., Maertens de Noordhout, A., Nakashima, K., Shannon, K., & Marsden, C. D. (1989). Delay in the execution of voluntary movement by electrical or magnetic brain stimulation in intact man. Evidence for the storage of motor programs in the brain. Brain: A Journal of Neurology, 112 ( *Pt 3*), 649–663. https://doi.org/10.1093/brain/112.3.649

Denyer, R., Greenhouse, I., & Boyd, L. A. (2023). PMd and action preparation: Bridging insights between TMS and single neuron research. Trends in Cognitive Sciences. https://doi.org/10.1016/j.tics.2023.05.001

Diedrichsen, J., Wiestler, T., & Krakauer, J. W. (2013). Two Distinct Ipsilateral Cortical Representations for Individuated Finger Movements. Cerebral Cortex, 23(6), 1362– 1377. https://doi.org/10.1093/cercor/bhs120

Duque, J., Labruna, L., Cazares, C., & Ivry, R. B. (2014). Dissociating the influence of response selection and task anticipation on corticospinal suppression during response preparation. Neuropsychologia, 65, 287–296. https://doi.org/10.1016/j.neuropsychologia.2014.08.006

Duque, J., Labruna, L., Verset, S., Olivier, E., & Ivry, R. B. (2012). Dissociating the role of prefrontal and premotor cortices in controlling inhibitory mechanisms during motor preparation. Journal of Neuroscience, 32(3), 806–816.

Fujiyama, H., Van Soom, J., Rens, G., Cuypers, K., Heise, K. F., Levin, O., & Swinnen, S. P. (2016). Performing two different actions simultaneously: The critical role of interhemispheric interactions during the preparation of bimanual movement. Cortex, 77, 141–154. https://doi.org/10.1016/j.cortex.2016.02.007

Ganguly, K., Secundo, L., Ranade, G., Orsborn, A., Chang, E. F., Dimitrov, D. F., Wallis, J. D., Barbaro, N. M., Knight, R. T., & Carmena, J. M. (2009). Cortical Representation of Ipsilateral Arm Movements in Monkey and Man. Journal of Neuroscience, 29(41), 12948–12956. https://doi.org/10.1523/JNEUROSCI.2471-09.2009

Godschalk, M., Lemon, R. N., Kuypers, H. G., & van der Steen, J. (1985). The involvement of monkey premotor cortex neurones in preparation of visually cued arm movements. Behavioural Brain Research, 18(2), 143–157. https://doi.org/10.1016/0166-4328(85)90070-1

Greenhouse, I., Sias, A., Labruna, L., & Ivry, R. B. (2015). Nonspecific Inhibition of the Motor System during Response Preparation. J Neurosci, 35(30), 10675–10684. https://doi.org/10.1523/JNEUROSCI.1436-15.2015

Groppa, S., Schlaak, B. H., Münchau, A., Werner-Petroll, N., Dünnweber, J., Bäumer, T., van Nuenen, B. F. L., & Siebner, H. R. (2012). The human dorsal premotor cortex facilitates the excitability of ipsilateral primary motor cortex via a short latency cortico-cortical route. Human Brain Mapping, 33(2), 419–430.

Groppa, S., Werner-Petroll, N., Münchau, A., Deuschl, G., Ruschworth, M. F. S., & Siebner, H. R. (2012). A novel dual-site transcranial magnetic stimulation paradigm to probe fast facilitatory inputs from ipsilateral dorsal premotor cortex to primary motor cortex. Neuroimage, 62(1), 500–509.

Hannah, R., Cavanagh, S. E., Tremblay, S., Simeoni, S., & Rothwell, J. C. (2018). Selective Suppression of Local Interneuron Circuits in Human Motor Cortex Contributes to Movement Preparation. Journal of Neuroscience, 38(5), 1264–1276. https://doi.org/10.1523/JNEUROSCI.2869-17.2017

Hayward, K. S., Neva, J. L., Mang, C. S., Peters, S., Wadden, K. P., Ferris, J. K., & Boyd, L. A. (2017). Interhemispheric pathways are important for motor outcome in individuals with chronic and severe upper limb impairment post stroke. Neural Plasticity, 2017.

Hocherman, S., & Wise, S. P. (1991). Effects of hand movement path on motor cortical activity in awake, behaving rhesus monkeys. Experimental Brain Research, 83(2), 285–302. https://doi.org/10.1007/BF00231153

Jackson, N., & Greenhouse, I. (2019). VETA: an open-source Matlab-based toolbox for the collection and analysis of electromyography combined with transcranial magnetic stimulation. Frontiers in Neuroscience, 13, 975.

Jenkinson, M., Beckmann, C. F., Behrens, T. E., Woolrich, M. W., & Smith, S. M. (2012). FSL. Neuroimage, 62(2), 782–790. https://doi.org/10.1016/j.neuroimage.2011.09.015

Kaufman, M. T., Churchland, M. M., Ryu, S. I., & Shenoy, K. V. (2014). Cortical activity in the null space: Permitting preparation without movement. Nature Neuroscience, 17(3), 440– 448.

Kaufman, M. T., Seely, J. S., Sussillo, D., Ryu, S. I., Shenoy, K. V., & Churchland, M. M. (2016). The largest response component in the motor cortex reflects movement timing but not movement type. Eneuro, 3(4).

Kazennikov, O., Hyland, B., Wicki, U., Perrig, S., Rouiller, E. M., & Wiesendanger, M. (1998). Effects of lesions in the mesial frontal cortex on bimanual co-ordination in monkeys. Neuroscience, 85(3), 703–716. https://doi.org/10.1016/s0306-4522(97)00693-3

Klein, P. A., Duque, J., Labruna, L., & Ivry, R. B. (2016). Comparison of the two cerebral hemispheres in inhibitory processes operative during movement preparation. Neuroimage, 125, 220–232. https://doi.org/10.1016/j.neuroimage.2015.10.007

Kleiner, M., Brainard, D., Pelli, D., Ingling, A., Murray, R., & Broussard, C. (2007). What’s new in psychtoolbox-3. Perception, 36(14), 1–16.

Koch, G., Franca, M., Del Olmo, M. F., Cheeran, B., Milton, R., Alvarez Sauco, M., & Rothwell, J. C. (2006). Time course of functional connectivity between dorsal premotor and contralateral motor cortex during movement selection. J Neurosci, 26(28), 7452–7459. https://doi.org/10.1523/JNEUROSCI.1158-06.2006

Kroeger, J., Bäumer, T., Jonas, M., Rothwell, J. C., Siebner, H. R., & Münchau, A. (2010). Charting the excitability of premotor to motor connections while withholding or initiating a selected movement. European Journal of Neuroscience, 32(10), 1771–1779.

Leocani, L., Cohen, L. G., Wassermann, E. M., Ikoma, K., & Hallett, M. (2000). Human corticospinal excitability evaluated with transcranial magnetic stimulation during different reaction time paradigms. Brain, 123 ( *Pt 6*), 1161–1173. https://doi.org/10.1093/brain/123.6.1161

Liuzzi, G., Hörniß, V., Hoppe, J., Heise, K., Zimerman, M., Gerloff, C., & Hummel, F. C. (2010). Distinct Temporospatial Interhemispheric Interactions in the Human Primary and Premotor Cortex during Movement Preparation. Cerebral Cortex, 20(6), 1323–1331. https://doi.org/10.1093/cercor/bhp196

Liuzzi, G., Hörniss, V., Zimerman, M., Gerloff, C., & Hummel, F. C. (2011). Coordination of uncoupled bimanual movements by strictly timed interhemispheric connectivity. J Neurosci, 31(25), 9111–9117. https://doi.org/10.1523/JNEUROSCI.0046-11.2011

Mars, R. B., Bestmann, S., Rothwell, J. C., & Haggard, P. (2007). Effects of motor preparation and spatial attention on corticospinal excitability in a delayed-response paradigm. Experimental Brain Research, 182(1), 125–129. https://doi.org/10.1007/s00221-007-1055-4

Merrick, C. M., Dixon, T. C., Breska, A., Lin, J., Chang, E. F., King-Stephens, D., Laxer, K. D., Weber, P. B., Carmena, J., Thomas Knight, R., & Ivry, R. B. (2022). Left hemisphere dominance for bilateral kinematic encoding in the human brain. ELife, 11, e69977. https://doi.org/10.7554/eLife.69977

Messier, J., & Kalaska, J. F. (2000). Covariation of primate dorsal premotor cell activity with direction and amplitude during a memorized-delay reaching task. J Neurophysiol, 84(1), 152–165. https://doi.org/10.1152/jn.2000.84.1.152

Mitsuhashi, M., Yamaguchi, R., Kawasaki, T., Ueno, S., Sun, Y., Isa, K., Takahashi, J., Kobayashi, K., Onoe, H., Takahashi, R., & Isa, T. (2023). State-dependent role of interhemispheric pathway for motor recovery in primates (p. 2023.04.23.538021). bioRxiv. https://doi.org/10.1101/2023.04.23.538021

Mochizuki, H., Huang, Y., & Rothwell, J. C. (2004). Interhemispheric interaction between human dorsal premotor and contralateral primary motor cortex. The Journal of Physiology, 561(1), 331–338.

Ni, Z., Gunraj, C., Nelson, A. J., Yeh, I. J., Castillo, G., Hoque, T., & Chen, R. (2009). Two phases of interhemispheric inhibition between motor related cortical areas and the primary motor cortex in human. Cereb Cortex, 19(7), 1654–1665. https://doi.org/10.1093/cercor/bhn201

Noskin, O., Krakauer, J. W., Lazar, R. M., Festa, J. R., Handy, C., O’Brien, K. A., & Marshall, R. S. (2008). Ipsilateral motor dysfunction from unilateral stroke: Implications for the functional neuroanatomy of hemiparesis. Journal of Neurology, Neurosurgery, and Psychiatry, 79(4), 401–406. https://doi.org/10.1136/jnnp.2007.118463

Oldfield, R. C. (1971). The assessment and analysis of handedness: The Edinburgh inventory. Neuropsychologia, 9(1), 97–113.

O’Shea, J., Sebastian, C., Boorman, E. D., Johansen-Berg, H., & Rushworth, M. F. S. (2007). Functional specificity of human premotor–motor cortical interactions during action selection. European Journal of Neuroscience, 26(7), 2085–2095.

Picard, N., & Strick, P. L. (2001). Imaging the premotor areas. Current Opinion in Neurobiology, 11(6), 663–672.

Riehle, A., & Requin, J. (1989). Monkey primary motor and premotor cortex: Single-cell activity related to prior information about direction and extent of an intended movement. J Neurophysiol, 61(3), 534–549. https://doi.org/10.1152/jn.1989.61.3.534

Rurak, B. K., Rodrigues, J. P., Power, B. D., Drummond, P. D., & Vallence, A. M. (2021a). Test Re-test Reliability of Dual-site TMS Measures of SMA-M1 Connectivity Differs Across Inter-stimulus Intervals in Younger and Older Adults. Neuroscience, 472, 11–24. https://doi.org/10.1016/j.neuroscience.2021.07.023

Rurak, B. K., Rodrigues, J. P., Power, B. D., Drummond, P. D., & Vallence, A.-M. (2021b). Reduced SMA-M1 connectivity in older than younger adults measured using dual-site TMS. European Journal of Neuroscience, 54(7), 6533–6552. https://doi.org/10.1111/ejn.15438

Serrien, D. J., Sovijärvi-Spapé, M. M., & Rana, G. (2012). Subliminal priming and effects of hand dominance. Acta Psychologica, 141(1), 73–77. https://doi.org/10.1016/j.actpsy.2012.07.008

Shen, Y.-C., & Franz, E. A. (2005). Hemispheric competition in left-handers on bimanual reaction time tasks. Journal of Motor Behavior, 37(1), 3–9. https://doi.org/10.3200/JMBR.37.1.3-9

Sussillo, D., Churchland, M. M., Kaufman, M. T., & Shenoy, K. V. (2015). A neural network that finds a naturalistic solution for the production of muscle activity. Nat Neurosci, 18(7), 1025–1033. https://doi.org/10.1038/nn.4042

Taniguchi, Y., Burle, B., Vidal, F., & Bonnet, M. (2001). Deficit in motor cortical activity for simultaneous bimanual responses. Experimental Brain Research, 137(3), 259–268. https://doi.org/10.1007/s002210000661

Thura, D., Cabana, J.-F., Feghaly, A., & Cisek, P. (2022). Integrated neural dynamics of sensorimotor decisions and actions. PLOS Biology, 20(12), e3001861. https://doi.org/10.1371/journal.pbio.3001861

Tokuno, H., & Nambu, A. (2000). Organization of nonprimary motor cortical inputs on pyramidal and nonpyramidal tract neurons of primary motor cortex: An electrophysiological study in the macaque monkey. Cerebral Cortex, 10(1), 58–68.

Verstraelen, S., van Dun, K., Depestele, S., Van Hoornweder, S., Jamil, A., Ghasemian-Shirvan, E., Nitsche, M. A., Van Malderen, S., Swinnen, S. P., Cuypers, K., & Meesen, R. L. J. (2021). Dissociating the causal role of left and right dorsal premotor cortices in planning and executing bimanual movements – A neuro-navigated rTMS study. Brain Stimulation, 14(2), 423–434. https://doi.org/10.1016/j.brs.2021.02.006

Wiesendanger, M., Wicki, U., & Rouiller, E. (1994). 9—Are There Unifying Structures in the Brain Responsible for Interlimb Coordination? In S. P. Swinnen, H. Heuer, J. Massion, & P. Casaer (Eds.), Interlimb Coordination (pp. 179–207). Academic Press. https://doi.org/10.1016/B978-0-12-679270-6.50014-0

Wright, D. J., Wood, G., Franklin, Z. C., Marshall, B., Riach, M., & Holmes, P. S. (2018). Directing visual attention during action observation modulates corticospinal excitability. PLoS ONE, 13(1), e0190165. https://doi.org/10.1371/journal.pone.0190165

